# Human cortical organoids expose a differential function of GSK3 on direct and indirect neurogenesis

**DOI:** 10.1101/484741

**Authors:** Alejandro López-Tobón, Carlo Emanuele Villa, Cristina Cheroni, Sebastiano Trattaro, Nicolò Caporale, Paola Conforti, Raffaele Iennaco, Maria Lachgar, Marco Tullio Rigoli, Berta Marcó de la Cruz, Pietro Lo Riso, Erika Tenderini, Flavia Troglio, Marco de Simone, Isabel Liste-Noya, Stefano Piccolo, Giuseppe Macino, Massimiliano Pagani, Elena Cattaneo, Giuseppe Testa

**Affiliations:** Laboratory of Stem Cell Epigenetics, IEO, European Institute of Oncology, IRCCS, Milan, Italy Milan, Italy; Department of Oncology and Hemato-oncology, Università degli studi di Milano, Milan, Italy; Istituto Nazionale Genetica Molecolare, INGM, Milan, Italy; Unidad de Regeneración Neural, Unidad Funcional de Investigación de Enfermedades Crónicas, Instituto de Salud Carlos III (ISCIII), Madrid, Spain; Istituto Nazionale Genetica Molecolare INGM ‘Romeo ed Enrica Invernizzi’, Milan 20122, Italy.; Department of Medical Biotechnology and Translational Medicine, Università degli Studi di Milano, Milano; Department of Cellular Biotechnologies and Hematology, La Sapienza University of Rome, Rome, Italy 20129, Italy.

**Keywords:** human brain organoids, single cell, GSK3, corticogenesis

## Abstract

The regulation of proliferation and polarity of neural progenitors is crucial for the development of the brain cortex, with modes and timings of cell division intimately related to the stereotypical acquisition of layer-specific neuronal identities. Animal studies have implicated glycogen synthase kinase 3 (GSK3) as a pivotal regulator of both proliferation and polarity, yet the functional relevance of its signaling for the unique features of human corticogenesis remain to be elucidated. Here we harness human cortical brain organoids to probe the longitudinal impact of GSK3 inhibition through multiple developmental stages. Our results indicate that chronic GSK3 inhibition increases the proliferation of neural progenitors and causes massive derangement of cortical tissue architecture. Surprisingly, single cell transcriptome profiling revealed only a discrete impact on early neurogenesis and uncovered a pivotal role of GSK3 in the regulation of NEUROD1/2 lineages and outer radial glia (oRG) output, without compromising direct neurogenic trajectories. Through this first single cell-level dissection of the GSK3 regulatory network in human corticogenesis, our work underscores the robustness of transcriptional programs in determining neuronal identity independent of tissue architecture.

## Introduction

Neurogenesis is initiated by the formation of a neuroepithelium, composed by neural stem cells (NSCs) organized in apico-basal orientation that divide symmetrically leading to a rapid expansion of the NSC pool (Taverna et al., 2014; Temple, 2003). The polarization of the neuroepithelium precedes the differentiation of NSCs into radial glia cells (RGCs), and triggers the elongation of cytoplasmic processes that form a migratory scaffold for newborn neurons (Betizeau et al., 2013). Such polarization is in turn necessary for the acquisition of the key properties that define tissue organization (Johansson et al., 2010).

Glycogen synthase kinase 3 alfa and beta (GSK3α and β) are serine/threonine kinases encoded by 2 different genes, which function as integrating hubs for multiple proliferation and differentiation signals due to their central role in the receptor tyrosine kinase, Wnt and sonic hedgehog signaling pathways (Kim et al., 2009; McCubrey et al., 2016). GSK3 is involved in neurodevelopment through the phosphorylation of a broad set of substrates, including transcription factors essential for brain development, such as CREB (Grimes and Jope, 2001a), neurogenin2 (Ma et al., 2008) β-catenin (Aberle et al., 1997) and multiple microtubule-associated proteins (Fumoto et al., 2008; Zhou et al., 2004). Studies in animal models have provided a wealth of evidence linking GSK3 activity to the regulation of early and late neurogenesis in a stage-wise fashion. Its activity is required to maintain the overall polarity of the radial glia scaffold (Yokota et al., 2010). Genetic ablation of both GSK3α or GSK3β in radial glial cells (RGCs) results in a massive increase in neural progenitor proliferation with marked suppression of intermediate progenitor cells (IPCs) and postmitotic neurons (Kim et al., 2009). However, paralog-specific knockdown at later stages results in distinct outcomes, with the loss of GSK3β markedly decreasing the production of IPCs and upper-layer Cux1-positive neurons (Ma et al., 2017).

Activation of Wnt signaling as a result of GSK3 inhibition has been implicated both in neuroepithelial and apical radial glial cell expansion as well as in differentiation of basal progenitors, with the final outcome largely depending on the signaling context (Harrison-Uy and Pleasure, 2012). The inhibition of GSK3 during neural induction of mouse pluripotent stem cells results in the generation of neural progenitors of different rostro-caudal identity in a dose-dependent manner, whereas higher levels of inhibitor induce cells with more caudal identity, such as the midbrain (Hirabayashi et al., 2004; Kuwahara et al., 2010). Likewise, canonical Wnt pathway is active in cortical progenitor cells during early forebrain development in a high-medial to low-lateral pattern and as development progresses, cortical Wnt signaling decreases (Backman et al., 2005).

Despite the abundant evidence connecting GSK3 to neurogenesis in various animal models, much less is known of its role in the far more complex human context, mostly due to the lack of models that efficiently recapitulate the human-specific range of progenitor subpopulations. Seminal work conducted in human embryonic stem cells (hESCs) demonstrated their proficiency to form 3-dimensional aggregates containing self-organized apico-basally polarized cortical tissues with neurogenic properties (Watanabe et al., 2005). These 3-dimensional aggregates are able to generate features usually absent in monolayer cultures such as specific progenitor subpopulations and organizing centers (Eiraku et al., 2008; Kadoshima et al., 2013), constituting the precursors of current brain organoids protocols. Brain organoids have emerged as the most promising alternative to model neurodevelopment under a strictly human genetic background (Di Lullo and Kriegstein, 2017). Their ontogeny recapitulates most of the salient features of early to mid-fetal brain development, including progenitor populations and distribution of cell domains (Lancaster et al., 2013; Paşca et al., 2015). Strikingly, brain organoids show transcriptional and epigenetic signatures, such as global patterns of non-CG methylation, unique to the human cortical development (Luo et al., 2016). Single cell transcriptional profiling of organoids revealed a cellular diversity that closely matches in composition and transcriptional landscape the human fetal brain and confirmed the presence of human progenitor populations responsible for neocortical expansion (Camp et al., 2015; Quadrato et al., 2017; Amiri et al., 2018).

Here we explore the role of GSK3 on early to middle corticogenesis through the chronic and specific inhibition of its activity in human cortical organoids. By combining morphological characterization with massive parallel RNA sequencing on bulk and single cells, we uncover the molecular pathways modulated by GSK3 and breakdown their effect on distinct cell subpopulations of the developing human cortex, revealing a differential impact on neuronal progenitor subtypes and oRG-dependent lineages.

## Materials and Methods

### hPSC culture

hPSC lines were cultured under feeder-free conditions on Matrigel (BD Biosciences) coated dishes (diluted 1:40 matrigel:DMEM/F12) and grown in Tesr™ E8™ medium (Stem Cell Technologies). Cells were passaged upon treatment with ReLeSR (Stem Cell Technologies). All differentiation procedures were performed on iPSC lines with at least 15 passages after reprogramming. Pluripotent lines came from different individuals representing either human iPSC or ESC lines previously described (Adamo et al., 2015; González et al., 2014). All the cultures are regularly tested and maintained mycoplasma free. The distribution of lines across experiments is delineated in Supplementary table 1.

### Neural induction in 2D and lumen quantification

Briefly, hPSC cells were plated at a density of 0.7 × 10^5^ cells cm^2^ on Matrigel (BD Biosciences) coated dishes in grow medium supplemented with 10 µM ROCK inhibitor (Y-2763221, Cell Guidance System). Cell cultures were expanded for two days until they were 70% confluent. The starting differentiation medium includes DMEM/F12 (Life Technologies) with N2 and B27 without retinoic acid (Life Technologies), supplemented with 500 nM LDN193189 (Sigma) and 10 µM SB431542 (Tocris). From day 0 CHIR99021 was added to the medium at 1 μM concentration in parallel with neural induction without CHIR99021 until day 20. At day 12 cells were moved to neural medium containing DMEM/F12 (Life Technologies) with N2 and B27 plus retinoic acid and 30ng ml-1 BDNF. For lumen quantification cells were fixed at day 20 in 4% (wt/vol) paraformaldehyde (PFA) for 15 minutes at room temperature (RT) and washed 3 times with phosphate-buffered saline (PBS). Cells were then permeabilized with PBS containing 0.5% Triton X-100 (Sigma) and blocked with 10% (vol/vol) normal goat serum (NGS; Vector) for 1 hour at RT. Next, cells were incubated overnight at 4°C. The following primary antibodies and dilutions were used: anti-Ncadherin, 1:800 (BD Biosciences) and anti-PALS1, 1:500 (Cell Signaling Technology). Object identification module of Cell Profiler software (v.2.1.1) was used to automatically quantify lumens (number and size of Pals1 positive areas) of neural rosettes.

### Proliferation assay

Number of viable proliferating cells was estimated by luminiscence assay CellTiter-Glo^®^ (Promega). Briefly, 2×10^3^ cells/well were plated in 96-well flat bottom plates (Corning) with 4 replicates per condition and left to proliferate for 96 hours. Measurements were performed every 24 hours. Each data point was normalized to a blank from unseeded wells.

### Cortical Organoid differentiation and inhibition

Cortical organoids were differentiated as previously described (Paşca et al., 2015). Detailed protocol in *supplemental experimental procedures*. Chronic GSK3 inhibition was performed by adding CHIR99021 (Merck SML1046) to the medium at day 0 (1 µM) and kept throughout the differentiation process until reaching the respective sample collection timepoints.

### Growth curve

Organoids were moved at day 0 to 96-well U-bottom ultra-low attachment plates (Corning) and kept individually in order to avoid fusions for image acquisition until day 12 of differentiation. From day 12 onwards, organoids were moved individually to 24-well ultra-low attachment plates (Corning). Images were acquired with an EVOS Cell imaging System XL (Thermo) at the indicated differentiation days. Organoid size was calculated using an in-house developed a custom-script (by CEV) for FIJI software (v.1.49 NIH-USA).

### Tissue preservation and staining

Organoids were fixed in 4% (vol/vol) paraformaldehyde (PFA) for a minimum of 2h for day 18 organoids to overnight for day 50. Fixed organoids were washed twice with PBS and mounted on OCT cryopreservation medium on dry ice. Cryoblocks were preserved at −80 °C until the moment of sectioning. Cryosections were prepared using Leica CM 1900 instrument with 5 µm thickness. Sections were incubated with 10 mM Sodium citrate buffer (Normapur) for 45 min at 95°C + Tween 20 0,5% for simultaneous antigen retrieval and permeabilization. Antibody incubation details and list in *supplemental experimental procedures.* Images were acquired with a Leica DMI 6000B microscope (5×, 10×, and 20× objectives) and analyzed with LAS-AF imaging software and then processed using Image J (v1.49 NIH, USA) to adjust contrast for optimal RGB rendering. Confocal images were acquired with Leica SP8 microscope in resonant mode with Fluotar VISIR 25x/0.95 WATER objective, multiple tiles were acquired with PMT sensor and reconstructed via LASX software.

### Total RNA extraction and sequencing

Total RNA was isolated with the RNeasy Micro Kit (Qiagen, Hilden, Germany) according to the manufacturer’s instructions. RNA was quantified with Nanodrop and then the integrity was evaluated with Agilent 2100 Bioanalyzer (only if the quality ratios were not optimal after Nanodrop analysis). TruSeq Stranded Total RNA LT Sample Prep Kit (Illumina) was used for library preparation starting from 500 ng of total RNA for each sample. Sequencing was performed with the Illumina NOVAseq 6000 platform, with an average depth of 35 million 50bp paired-end reads per sample.

### Bulk Transcriptome analysis

*Differential expression analysis.* Three biological replicates were analyzed for untreated and treated organoids at each time point, for a total of 18 samples subjected to bulk RNA sequencing. Gene expression quantification at the gene level was performed by Salmon (version 0.8.2)(Patro et al., 2017), using hg38 RefSeq annotation. To estimate differential expression, the matrix of gene counts was analyzed by edgeR (version 3.20.9) (Robinson et al., 2009). For each time point, genes with an expression level of at least 2 cpm (count per million) in at least 3 samples were selected for the analysis. Small genes, ribosomal genes and fusion genes were excluded. After TMM normalization, differential expression analysis comparing treated to untreated samples was performed using a likelihood ratio test on the coefficients of a negative binomial model. Significantly modulated genes were selected setting an absolute value of log2 fold change (Log2FC) higher than 1 and a false discovery rate (FDR) lower than 5%. Log2 cpm values, were used for heatmap representation of gene expression profiles (visualized as z-scores). Heatmaps were produced with pheatmap R package (version 1.0.10, Raivo Kolde (2018). pheatmap: Pretty Heatmaps.). Analyses were performed in R version 3.4.4. Functional annotation of biological functions was performed by Gene ontology analysis and Gene set enrichment analysis (GSEA) using as set source H1 collection from the Molecular Signature Database (Liberzon et al., 2015). Detailed in *supplemental experimental procedures*.

### Single-Cell suspension, cDNA synthesis, library preparation and sequencing

Organoids were collected at day 50 or 100. 3-5 organoids per condition were dissociated by incubation with a solution of 0.5 mg/ml trypsin + 0,22 mg/ml EDTA (Euroclone) with 10 µl of DNaseI 1000 U/ml (Zymo Research) for 30 - 45 min according to organoid size. Digested suspensions were passed once through 0.4 µm Flowmi™ cell strainers, resuspended in PBS and counted using TC20 automatic cell counter (Biorad). Droplet-based single-cell partitioning and single-cell RNA-Seq libraries were generated using the Chromium Single-Cell 3′ Reagent v2 Kit (10× Genomics, Pleasanton, CA) following manufacturer’s instructions (Zheng et al., 2017) Detailed protocol in *Supplemental experimental procedures*.

### Single-Cell transcriptome analysis

Eleven biological samples (day 50: 3 untreated and 2 treated; day 100: 4 untreated and 2 treated) were examined by single cell analysis, for a total of 33293 cells and a median of 1733 features for cell. Libraries from single cell sequencing were aligned relying on the CellRanger v2.1 pipeline and using hg38 as reference. Before downstream analyses, data deriving from the 11 samples was integrated by Seurat v3.0-alpha analytical framework (Stuart et al., 2018). After normalization, anchors for data integration were identified considering 3000 anchor points (genes) and 40 dimensions. For data reduction, UMAP was applied with 50 nearest neighbors (nn); cluster initial positions were set considering PAGA node position (Scanpy v1.3.1) (Wolf et al., 2018). On the integrated dataset, clusters were identified by applying Louvain with Multilevel Refinement from Seurat with resolution parameter at 0.7. This resulted in the identification of 15 clusters. For cluster annotation, we applied for each of them the FindMarker Seurat function, using MAST as test and filtering for up-regulated genes with fold change > 1.5 and adjusted P value < 0.05. The obtained lists were compared in an overlap analysis with gene lists deriving from a fetal single cell datasets (https://cells.ucsc.edu/?ds=cortex-dev) (Amiri et al., 2018; Nowakowski et al., 2017). Cluster-specific expression levels of biologically-relevant genes identified among the top dysregulated were visualized by violin plots, stratified for stage and treatment. Cell cycle analysis were performed using Scanpy function score_genes_cell_cycle, relying on the genes from (Kowalczyk et al., 2015). Diffusion map algorithm for dimensionality reduction was performed with Scanpy with 50 nn. Pseudotime analysis for lineage branching reconstruction was applied using wishbone algorithm (Setty et al., 2016). The analysis was performed on the complete dataset, as well as separately for each of the four biological conditions in order to infer stage or treatment-selective trajectories; the origin was identified with the same method applied on complete dataset. Partition-based graph abstraction (PAGA) algorithm was applied on the complete dataset, as well as separately for each of the four biological conditions and plotted with layout Reingold Tilford. The position of the nodes identified on the complete dataset was exploited in the graph for each biological condition.

### Statistical Analysis

Statistical analyses were done using PRISM (GraphPad, version 6.0). Statistical significance was tested with the unpaired t-test, considering each line as biological replicates and comparing treatments as variables. N, p-values and significance are reported in each figure and legend. All results were expressed as means ± SD. No data points were excluded from the reported analyses.

## Results

### GSK3 is essential for cortical organoid morphogenesis

Patterned cortical organoids follow a stereotypical morphogenesis beginning with RGCs aligned in three-dimensions around ventricle-like structures (VLS), mimicking the hierarchical organization of ventricular zone of the dorsal telencephalon. Polarized RGCs are evident by day 18 (Figure 1A), with NESTIN+/PAX6 + cells comprising about 80% of the population (Figure 1B). This proportion decreases over time and becomes restricted to well-confined proliferative domains that gradually generate the neurons of the cortical plate (Paşca et al., 2015). In order to investigate the role of GSK3 activity throughout corticogenesis, we chronically exposed cortical organoids to the most specific GSK3 inhibitor available CHIR99021 (termed CHIR hereafter) at a 1 µM concentration, selected below the threshold for endodermal or mesodermal lineage induction in hPSCs (Patsch et al., 2015). Chronic GSK3 inhibition resulted in an increase in organoid size (Figures 1A, 2F) concomitant with a virtually complete loss of VLS formation (Figure 1A-B), an effect that became even more dramatic by day 50 (Figure. 1C-E). Unexpectedly, this difference in organoid size and radial organization was accompanied by a marginal decrease of PAX6+ cells at day 18 (Figure 1F) (CTL 76 ± 3.3, CHIR 65 ± 2.7), as well as of its expression levels, (Figure. S1A), while no significant difference was observed at day 50 in the proportion of PAX6+ cells (Figure 1G)(CTL 20 ± 1.2, CHIR 22 ± 1.0). Moreover, staining of early neuron markers TBR1 (layer V-VI neuronal progenitors) (Figure 1E) and DCX (Figure 1D) revealed a profound disarray in tissue architecture in the face of only a slight reduction in TBR1+ cells (CTL 39 ± 1.3, CHIR 32 ± 1.3) (Figure 1J), indicating that GSK3 activity is not essential for attaining early neurogenesis, but critical for the correct morphogenesis of the developing cortex.

**Figure 1.**
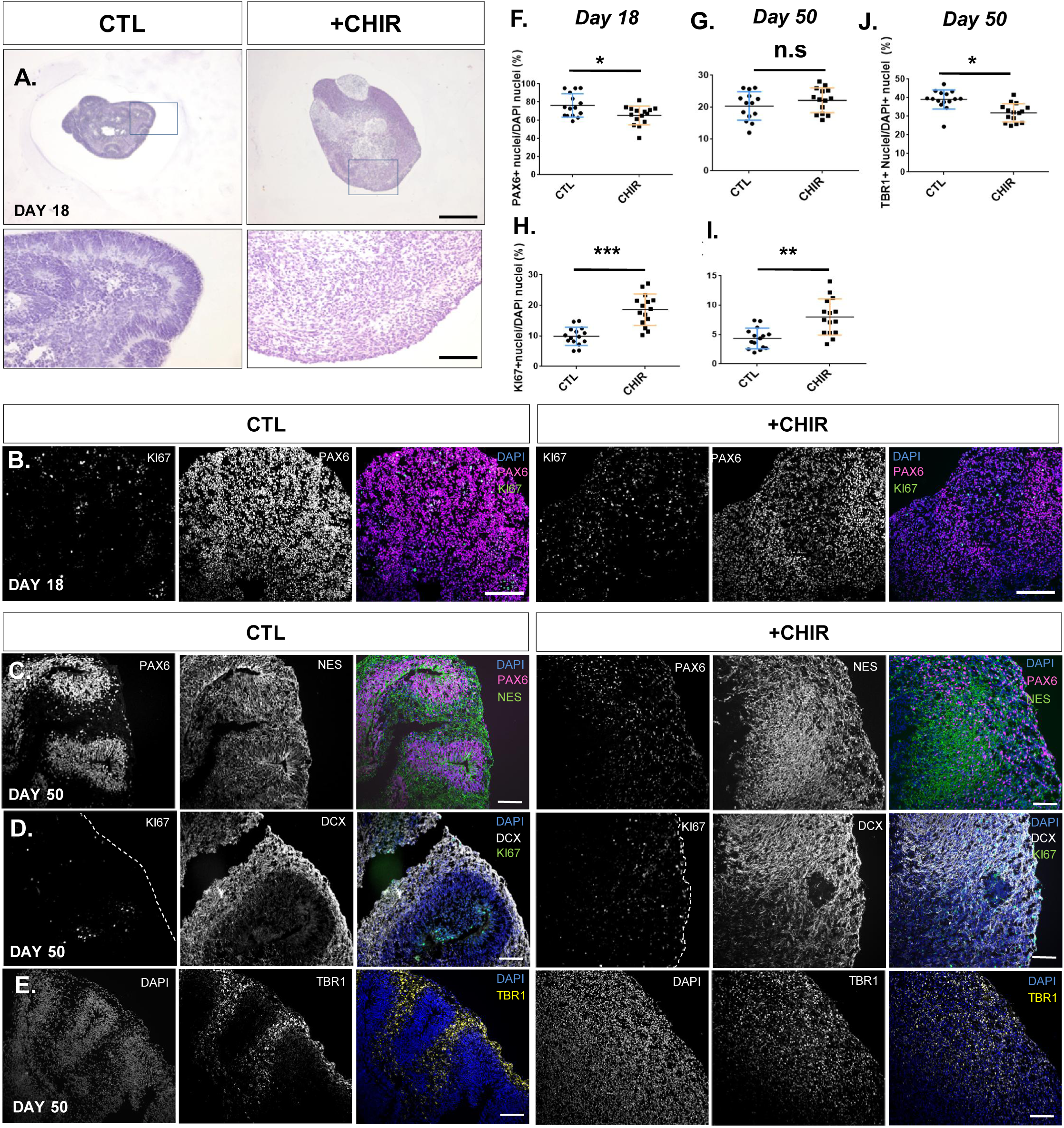
Morphogenetic alterations caused by chronic GSK3 inhibition. **A.** Representative captions of cortical organoids at 18 d day, stained with hematoxylin-eosin (upper - 10x, lower - 63x magnifications), scale bars 200, and 10 μm respectively. **B.** Representative images from day 18 organoids immunostained with anti-PAX6 (red), anti-Ki67 (green), DAPI (blue), widefield fluorescence images, scale bar = 50 um. **C-E.** Representative images from day 50 organoids immunostained with: **C.** anti-PAX6 (red), anti-Nestin (green), DAPI (blue), **D.** anti-DCX (white), anti-KI67 (green), DAPI (blue), **E** anti-TBR1 (yellow), DAPI (blue), scale bar = 50 μm**. F-G** Quantification of the proportion of PAX6+ nuclei relative to total nuclei (DAPI) at **F.** day 18 and **G.** day 50 of differentiation. **H-I.** Quantification of the proportion of Ki67+ nuclei relative to total nuclei (DAPI) at **H.** day 18 and **I.** day 50 of differentiation. **J.** Quantification of the proportion of TBR1+ nuclei relative to total nuclei (DAPI) in day 50 organoids. All quantifications were performed in 5 organoids per line, 3 independent hPSC lines (N = 15, n = 3), unpaired t-test *p<0.05, **p<0.01 and ***p<0.001, bars = mean + SD.

To dissect the mechanisms underlying this morphogenic defect, we adopted a two-tiered strategy: i) a validation in a classic 2D model attuned to quantify essential properties of neural stem cells such as polarity and proliferation (Conforti et al., 2018); and ii) transcriptional profiling of patterned cortical organoids at 50 and 100 days for a dynamic characterization of early corticogenesis (Figure. 2A). We followed the emergence of 2D neural rosettes until 20 days *in vitro*, when rosettes are typically PASL1+ at the apical end. GSK3 inhibition drastically reduced rosette number (Figure 2B-C) as well as the average size of formed rosettes (Figure 2D). In addition, quantification of the population size at the first 72 hours of dual-Smad inhibition revealed an increase in proliferation rate up to 1.5-fold compared to control (Figure 2E), indicating that both the morphogenic disarray and the increased size observed in early stage cortical organoids arise from aberrations already present during the transition from pluripotency to NSCs. We then validated these findings by quantifying the impact of GSK3 inhibition on organoid size through a growth measurement over a 40 days’ time-course. We found a sustained increase in organoid size upon chronic GSK3 inhibition (Figure 2F-H) with a rate surge that dramatically picks up in pace between days 10 and 15, subsequent to the time point when cortical organoids begin to be exposed to proliferation boosting factors FGF2 and EGF (Figure 2H), in agreement with the increased proliferation rate observed at the first 72 hours of neural induction (Figure. 2E) and pointing to a growth factor-mediated proliferation process.

**Figure 2.**
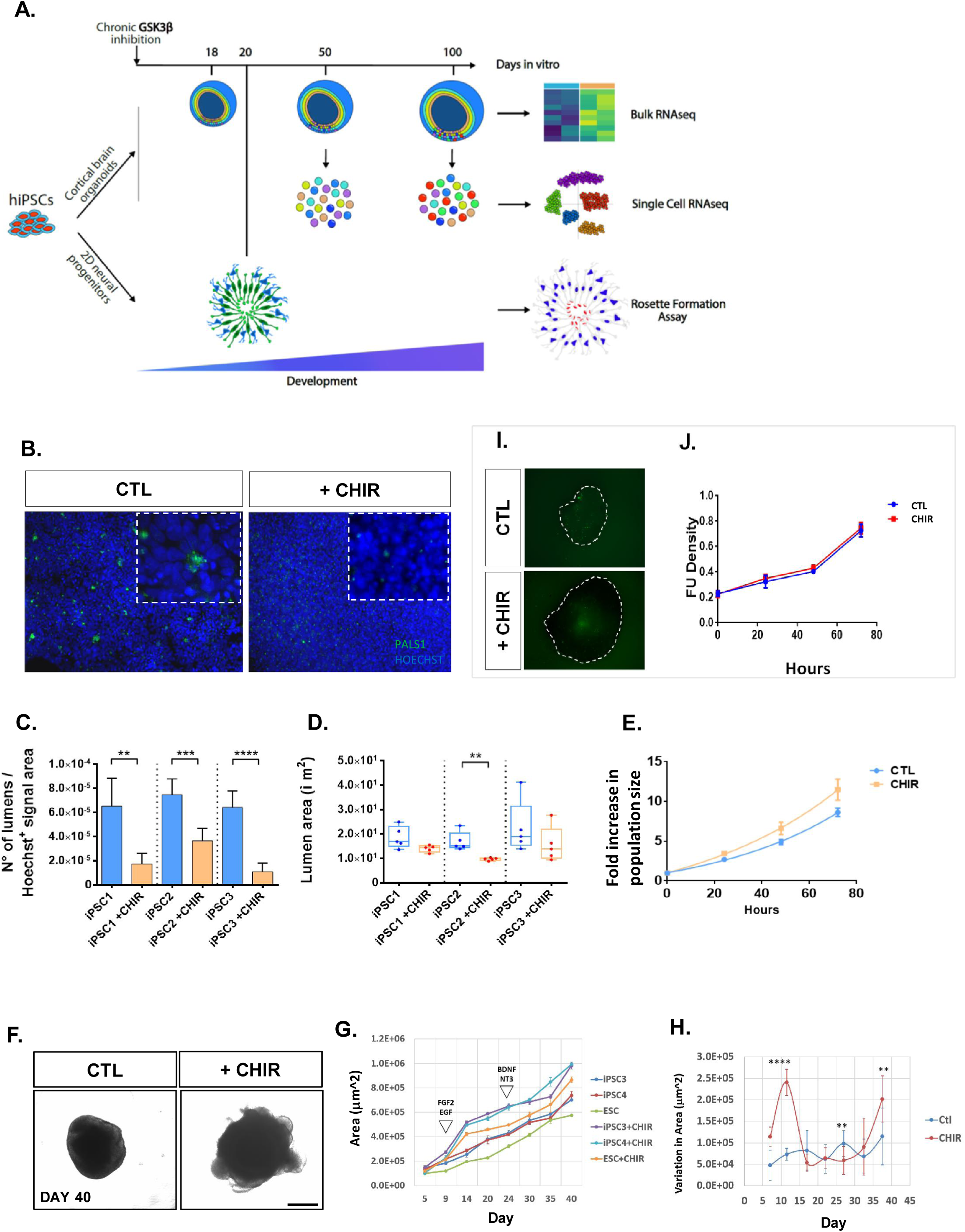
GSK3 inhibition disrupts lumen organization and progenitor proliferation rate. **A.** Experimental design: hPSCs differentiated following two parallel protocols, in 3D (up) cortical organoids or 2D (down) dual-smad inhibition. In both cases, parallel rounds were either exposed or not to GSK3 inhibitor CHI99021 (1 uM) starting from day 0 until indicated sample collection timepoint. **B.** Representative captions from immunostaining performed for anti-Pals1(green)/Hoechst(blue), widefield fluorescence images acquisition 20X, scale bar = 20 um. **C.** Bar-plots represent the average ± st.dev. of 5 independent images for lines/conditions; unpaired t-test **p<0.01, ***p<0.001 and ****p<0.0001. **D.** Lumen quantification (number and area) was performed by CellProfiler software. **E.** Cell proliferation rate was estimate by Cell titer Glo luminescence assay, measurements every with measurements every 24 hours for 96 hours in triplicate n = 3. **F.** Representative brightfield captions of day 40 organoids, scale bar = 1 mm. **G.** Growth curve performed in organoids differentiated for 40 days. Brightfield captions were taken at days 5, 9, 14, 20, 24, 30, 35, 40. Points represent the average of 4 organoids per line. Size quantifications were performed with a custom-made FIJI function. **H.** Derivative of the growth-rate delta between timepoints. The delta was computed on the average of untreated or treated samples. SD was calculated as a cumulative SD across all replicates and samples from each condition (n=9 per condition), unpaired t-test **p<0.01 and ****p<0.0001. **I.** Representative images from day 50 organoids stained with CellTox ® green as a marker of cytotoxicity after 72 hours of growth factor starvation (depletion B27 and growth factors). **J.** Quantification of CellTox ® fluorescence at 0, 24, 48 and 72 hours after growth factor starvation.

A previous report suggested that the sustained application of CHIR99021 in a patterned organoid protocol starting at day 12 of differentiation decreased the number of Caspase 3 + core cells (Qian et al., 2016), which could explain the increase in organoid size. To test whether chronic CHIR application had an overall effect on cell viability, we performed luminescent quantitation of membrane permeability in day 50 organoids every 24 hours, during 72 hours of growth factor starvation. Quantification showed no changes in cell death as a result of GSK3 inhibition, either in base line (0 h) or after 72 h of starvation (Figure 2I-J), showing that chronic GSK3 has no impact on basal or starvation-mediated cell death, hence excluding reduced apoptosis as a plausible cause of the increased organoid size. Instead, quantification of the proliferation marker KI67 showed a marked increase of actively dividing cells in organoids at day 18 (CTL 10 ± 0.7, CHIR 19 ± 1.3) (Figure 1H) and day 50 (CTL 4.3 ± 0.4, CHIR 8.0 ± 0.8) (Figure 1I) pointing to the regulation of progenitor proliferation as a core aspect of GSK.

### Transcriptional regulation associated to GSK3 activity throughout cortical development

Abundant evidence supports an active role of GSK3 in neurogenesis (Hur and Zhou, 2010), however, since complete GSK3 ablation causes systemic failure and early embryonic lethality (Hoeflich et al., 2000), its levels/activity are usually manipulated both *in vitro* and *in vivo* after the early stages of neurodevelopment have already been completed. The surprisingly mild effect of chronic GSK3 inhibition on neural maturation prompted us to dissect its role on gene expression by bulk RNAseq at three critical time-points recapitulating relevant stages of human cortical development (i.e. day 18: abundant apical radial glia; day 50: presence of intermediate progenitors and early neurons; and day 100: presence of lower layer neurons and beginning of astrogenesis) (Paşca et al., 2015).

Principal component analysis (PCA) indicated the stage of differentiation as the main source of variability and revealed an apparent delay of CHIR-treated organoids evident at days 50 and 100 (Figure. 3A). Principal component 2 revealed another aspect of CHIR treatment effect, which was strongly magnified at day 50 (Figure 3A). Analysis of the expression of gene signatures defining specific stages of differentiation (active proliferation, early and late RGs, early and late neurons) (Figure S1) confirmed a sustained increase of the proliferation marker MKI67 at all stages (Figure S1A-C), along with a differential impact on genes controlling distinct aspects of cell cycle progression at day 50 and day 100. Notably, we observed a robust upregulation of the anaphase promoting complex/cyclosome (APC/C), CDC20 (Figure S1B-C), which has been implicated in the modulation of NEUROD2 levels through the regulation of its ubiquitination (Yang et al., 2009). Interestingly, we observed a strong downregulation of NEUROD2 expression at day 50 and day 100 (Figure S1B-C) arguing for the existence of a regulatory feedback between CDC20-NEUROD2 at the transcriptional level, which may in turn result in the specific depletion of early-onset neuronal lineages (Telley et al., 2016). Moreover, the expression levels of the canonical neuronal maturity markers DCX, TUBB3, MAP2, MEF2C and STMN2 showed no generalized delay in maturation, but rather a differential modulation upon CHIR inhibition (exemplified by a contrasting upregulation of TUBB3 and downregulation MAP2) (Figure S1B-C), potentially resulting in an aberrant maturation process that could explain the downregulation of several AMPAR subunits observed at day 100 (Figure S1C-D)

**Figure 3.**
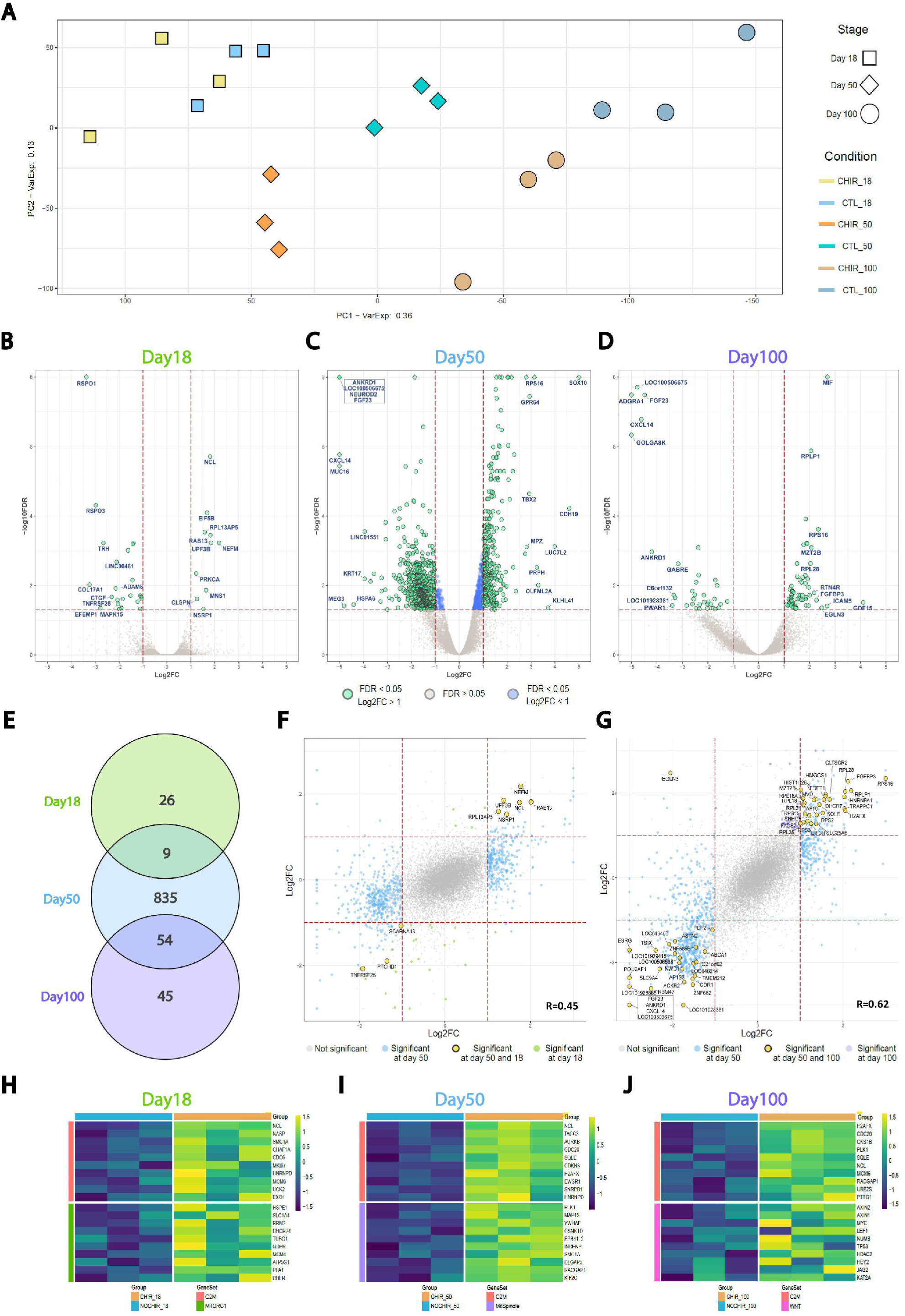
Impact of GSK3 inhibition on the transcriptional landscape. **A.** Principal Component Analysis performed on the whole transcriptome of untreated and CHIR-treated brain organoids at three stages of development: day 18, day 50 and day 100. **B-D.** Volcano plots illustrating the differential expression analysis for CHIR treatment at day 18 (B), day 50 (C) and day 100 (D). Results are reported for the pool of tested genes as −log10FDR and log2FC. Genes identified as significantly modulated (FDR < 5% and absolute log2FC > 1) are shown in green, while those respecting only the FDR threshold are depicted in blue and not significant genes (FDR > 5%) in grey. Gene symbols highlight the top-10 up-regulated and down-regulated genes (ranked by fold-change) in each stage. **E.** Venn diagram depicting the overlap of modulated genes across developmental stages. **F-G.** Scatterplots representing the relationship of the fold-change induced by GSK3 inhibition at day 50 and day18 (F) or day50 and day100 (G), for the subset of genes tested in both conditions. DEGs shared between the two examined conditions are reported in yellow, while DEGs specific of day 18, 50 or 100 in green, blue or violet respectively. Correlation coefficient calculated according the Spearman metrics. **H-J.** Gene expression profiles for selected genes sets significantly associated with CHIR treatment by GSEA (H: day 18; I: day 50; J: day 100). Expression levels (as z-score) for the 10 top-ranking genes in the leading edge are visualized for each gene set.

In order to address the functional implications of chronic CHIR exposure, we performed stage-wise differential expression analysis (DEA) (Figure 3, B-D). DEA confirmed the amplification of CHIR effect at day 50, with 898 differentially expressed genes (DEGs) (Figure 3C) (full list of DEGs, Supp. table 2). Ontology analysis revealed a persistent up-regulation of categories linked to cell proliferation and DNA replication (nucleosome assembly) and a down-regulation of cell surface components, as well as ion channels, including AMPA subunits (Figure S2B-C). While there were no genes differentially regulated upon CHIR treatment preserved at all stages, we identified 9 DEGs consistently dysregulated in the d18->d50 transition and 54 DEGs between day50->day100 (Figure 3E). Interestingly, several of the d18->d50 shared genes are linked to early neurogenesis and neuronal function (Figure 3F), while at d50->d100 to axon development and protein translation (Figure 3G). The overall comparison of gene modulation across stages showed a preserved trend of fold changes which was surprisingly stronger between day 50 -> day 100 (R = 0.62) compared to day 18 -> day 50 (R = 0.45), despite the longer timespan and increased cell-type diversity (Figure 3F,G), suggesting a more robust dependency of GSK3 activity occurring during this transition.

To gain deeper insight into the biological meaning of the coordinated modulation of functionally-related genes induced by GSK3 inhibition, we performed gene set enrichment analysis (GSEA). This approach confirmed a robust enrichment for pathways targeted by GSK3 activity (MTORC1, Myc signalling) and furthermore narrowed down the upregulation of G2-M transition modulators as a persistent feature of GSK3 transcriptional impact at all stages (Figure 3H-J). Likewise, several of the identified GSEA categories (complete list in Supp. table 3) are directly linked to the observed phenotypes, including the increased progenitor production (mTORC1 pathway), changes of cell polarity (mitotic spindle orientation) and neuronal fate (Wnt - β-Catenin pathway). Interestingly, both GO analysis at day 50 and GSEA (at all stages) point to the downregulation of inflammatory pathways by GSK3 inhibition, in line with a recognized role of GSK3 as mediator of neuroinflammation (Grimes and Jope, 2001b).

### Cortical organoids recapitulate the main features of mid-fetal human corticogenesis at the single cell level

Given that the transcriptional impact of GSK3 is greater at the day 50 -> day 100 transition, we harnessed single cell transcriptomics combined with distance-based analytical tools to break-down the effects of CHIR exposure in terms of the trajectories followed by specific cell populations during this developmental time frame. We carried out droplet-based single-cell mRNA sequencing to profile over 30000 cells (N =33293) in 11 biological samples from unexposed and exposed cortical organoids at day 50 and day 100 of differentiation. Projection of the expression levels of canonical population markers over UMAP, combined with the Louvain modularity algorithm (Šubelj and Bajec, 2011), identified 15 unsupervised clusters, which could be allocated into cell identities including early neurons (*DCX, ENO2*), postmitotic neurons (*STMN2*) and actively proliferating progenitors (*CDC20, MKI67*) (Figure 4A), bridged by areas rich in intermediate progenitors (*SMOC1*, *S100B*) and outer radial glia (oRG) (*HOPX-FAM107A*). By combining genetic signatures from single-cell RNA-seq studies of fetal human brain samples and cortical organoids (Amiri et al., 2018; Nowakowski et al., 2017), we were able to annotate clusters grouped into five *bona fide* population identities, radial glia progenitors (RGs), choroid, intermediate progenitors (IPCs), early neurons (EN) and late neurons (LN) (Figure 4B, Figure S3). We then investigated the underlying developmental trajectories using diffusion map algorithm (Coifman et al., 2005)(Figure 4C-D). Diffusion map confirmed the existence of one main developmental trajectory connecting RGs to postmitotic neurons linked by intermediate populations (Figure 4C). The application of pseudotime for lineage branching reconstruction (Haghverdi et al., 2016; Setty et al., 2016), with an origin anchored in the early progenitor compartment, unbiasedly reproduced the organization of populations from RG to LN (Figure 4D), which was also confirmed when repeated with an origin defined by different progenitor markers (data not shown). Likewise, the distribution of expression of specific population markers confirmed the presence of a lineage path from early proliferative populations through the intermediate progenitor-oRG transition and into early and late neural maturation stages (Figure 4E; MKI67 and CDK1 for proliferation; SMOC1 for intermediate progenitors; HOPX for oRG; DCX and STMN2 for neuronal maturity). Together, these results confirm that patterned cortical organoids recapitulate cardinal features of human corticogenesis including, the presence of IPCs, the emergence of oRG and its central position in neurogenic trajectories.

**Figure 4.**
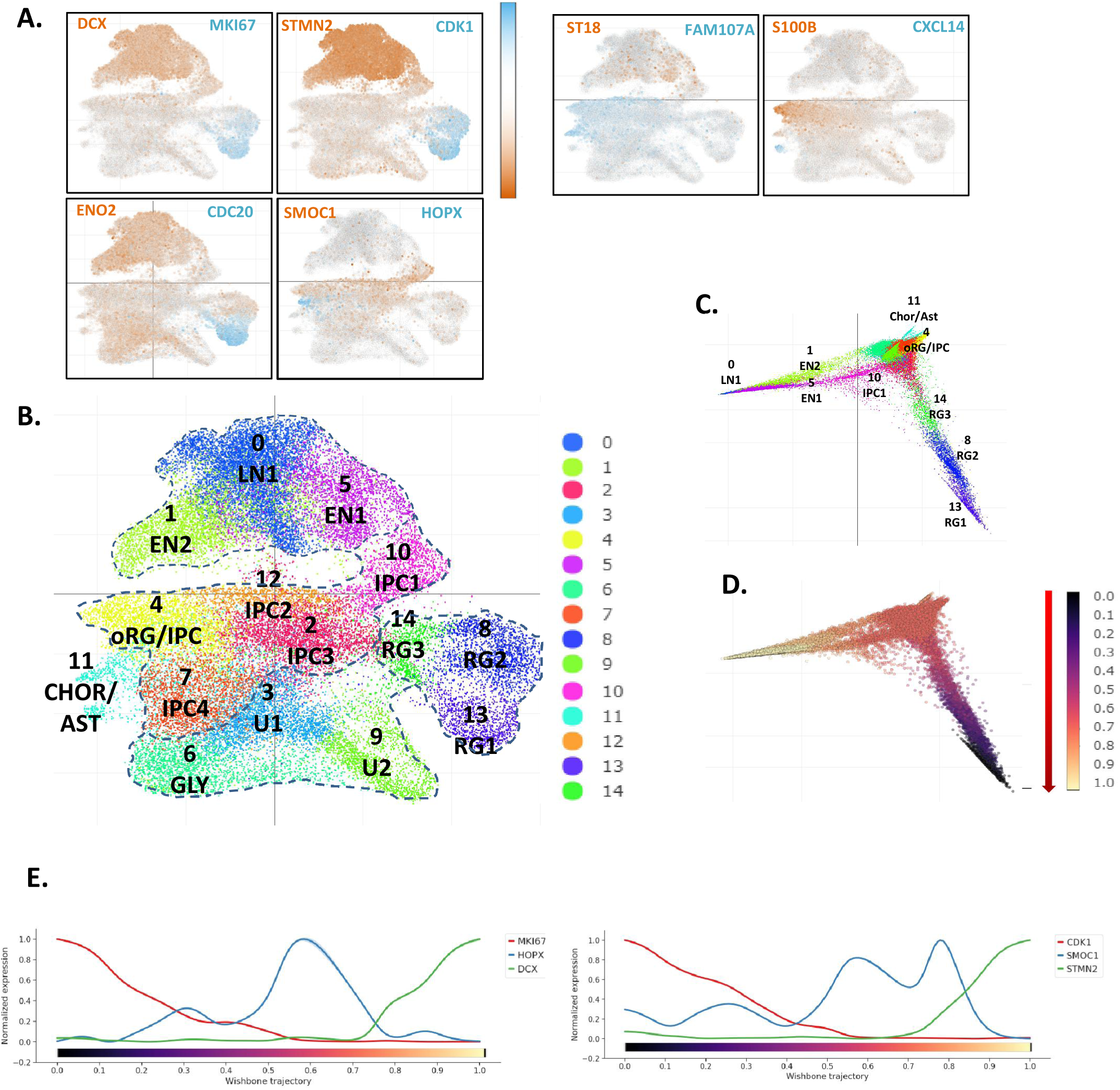
Cortical organoids recapitulate the main features of cortical development. **A.** UMAP plots. For each sub-panel, cells (represented as dots) are colored according to the expression levels of paired combinations of representative cell type markers (DCX, STMN2 = mature neurons), (ENO2, ST18 = early neural progenitors), (FAM107A, HOPX = outer radial glia), (SMOC1, S100B = intermediate progenitors) (CDK1, CDC20, MKI67 = proliferating progenitors) (CXCL14 = choroid/Astro). **B.** Louvain clusters in UMAP plot colored by identity; lines depict population areas defined by contrast with markers obtained from human fetal brain dataset (RG = Radial glia, IPC = Intermediate progenitor cells, EN = early neurons, LN = Late neurons, GLY = Glycolysis, CHOR/AST = choroid, astrocytes, U = unidentified). **C-D.** Diffusion map representing the developmental trajectory of the system. Cells (dots) are colored according to cluster identity (C) and to pseudotime trajectory (D), from origin in black to terminal state in light yellow according to wishbone algorithm. **E.** Visualization of the expression levels of representative genes along pseudotime, (CDK1, MKI67 = proliferative progenitors), (DCX, STMN2 = mature neurons), (HOPX = outer radial glia) (SMOC1 = intermediate progenitors).

### GSK3 inhibition differentially affects specific domains of corticogenesis

A comparative analysis of subpopulations revealed a selective impact of GSK3 inhibition on the relative proportion of specific cell subtypes (Figure 5A-E). A salient effect was the complete loss of a subpopulation characterized by the expression of a set of genes characteristic of choroid cells, (*CXCL14, HTR2C, TPD52L1, PCP4, EMX2*) (Figure 5A) (Nowakowski et al., 2017), in agreement with the strong downregulation of CXCL14 observed in bulk RNAseq at both day 50 and 100 (Figure 3B-D). Moreover, we observed a strong and specific depletion of NEUROD1, NEUROD2-at day 100 in neuronal clusters (0,1,5) without affecting NEUROG2 or NEUROD6 expressing cells (Figure 5B-C), underscoring the cardinal selectivity of GSK activity on these neurogenic pathways.

**Figure 5.**
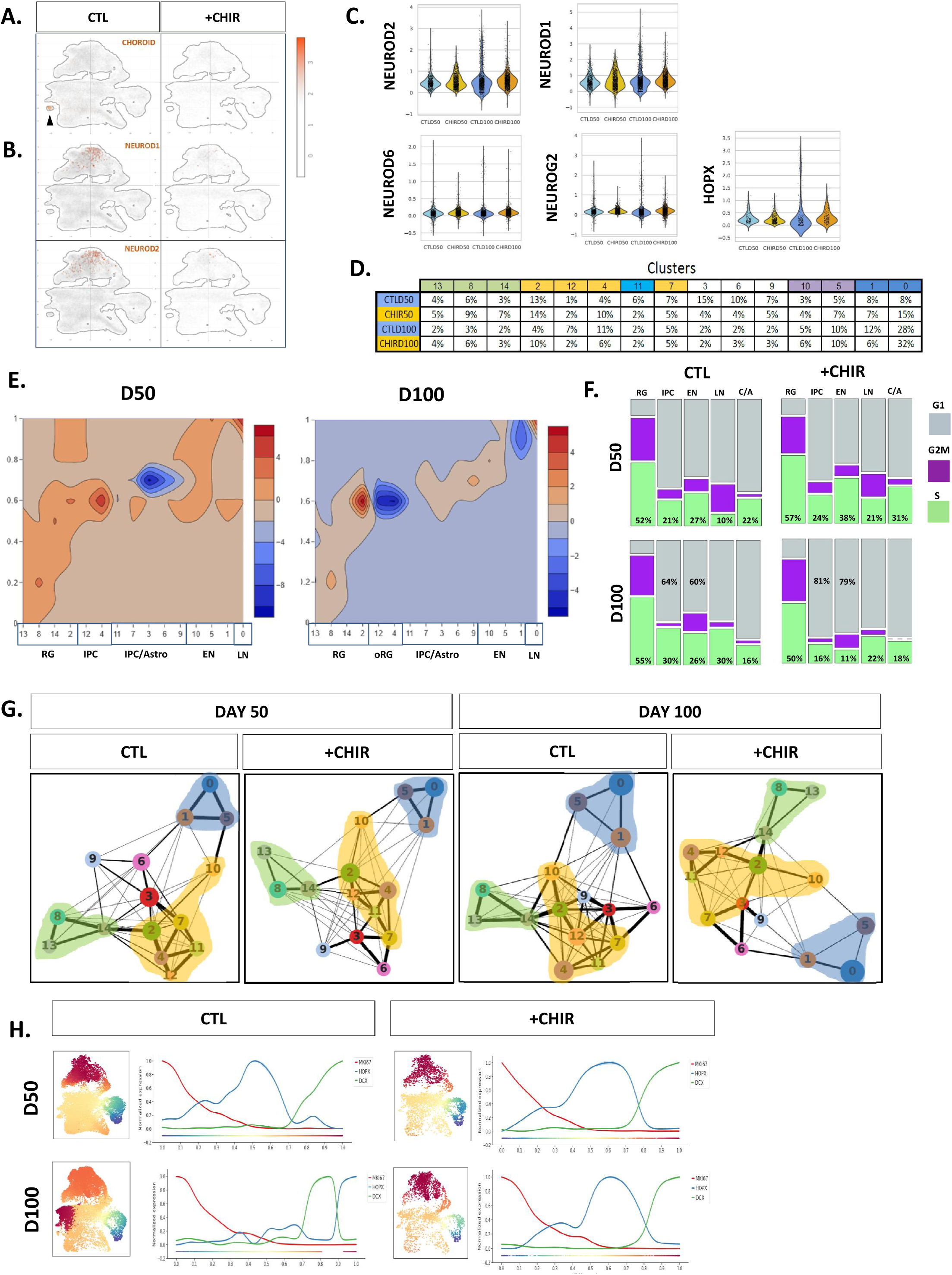
Effects of GSK3 inhibition at a single cell level in day 50 and day 100 cortical organoids. **A.** Z-score calculated on the expression levels of CXCL14, HTR2C, TPD52L1 and visualized as color-code on the UMAP identifies choroid cells. **B.** Visualization normalized expression levels of NEUROD2 or NEUROD1 as color code. **C.** Normalized expression levels and distribution of neural markers NEUROD2, NEUROD6 and NEUROG2 from cells assigned to clusters 0, 1 and 5 and outer radial glia marker HOPX from cells assigned to cluster 4. Levels are shown stratified by condition and timepoint; sub-sampling has been applied in order to visualize the same number of cells for each category. D. Distribution of cells (as percentage) in each cluster, stratified for time-point and condition. **E.** Contour plot representing the difference in frequency distribution in clusters and pseudotime intervals of treated versus untreated cells at day 50 and day 100. Red depicts values higher in CHIR-treated, while blue values higher in CTL. **F.** Distribution of cells across G1, S, G2/M phases of the cell cycle, divided by treatment and stage, in the following populations: apical progenitors (RG), intermediate progenitors (IPC), early maturity (EN), neural maturity (LN), astrogenesis (C/A). The mosaic plots report for each population the percentage of cells in each phase (G1:grey; S: green; G2M: purple). **G.** PAGA analysis applied stage and treatment-wise on the 15 clusters identified on the complete dataset. Circle diameter represents the fraction of cells assigned to each cluster; edge thickness visualizes the strength of connections across cells of the related clusters. Shadows highlight areas: proliferation (green), maturity (blue), intermediate (yellow). **H.** Over-imposition of pseudotime analyses performed separately for each experimental condition as color-scale on UMAP calculated on the complete system. Blue (origin); dark red (terminal state) according to wishbone trajectories. Visualization of the expression levels of representative genes along the condition-specific pseudotime.

Cluster-wise differential expression analysis, revealed selective effects on key genes influencing fate specification of postmitotic neurons (cluster 0) (*GRIA2, MAP2, BCL11A, PCDH9*), intermediate progenitors and outer radial glia (clusters 4,7) (*HES1, FABP5, HOPX, MT3, SPARCL1*), (Figure S4). Interestingly, the analysis of cell type distribution across pseudotime (Figure 5E) uncovered a higher proportion of cells in both progenitor and mature areas upon chronic GSK3 inhibition at day 50. This result was compatible with two possibilities: i) the cells identified as mature by predefined gene signatures could actually turn out to display mixed identities, co-expressing both progenitor and maturation markers as a result of a fundamental derangement of developmental hierarchies; or ii) the highly proliferative progenitors from CHIR-treated organoids favor direct over indirect transitions through the intermediate stages or even skip them altogether. In order to test these hypotheses, we plotted multiple proliferating/maturity signatures into UMAP and found no evidence of overlap in CHIR-treated organoids (Figure S5A); likewise, we did not observe an overlapping distribution between ki67 + and DCX+ cells (Figure 1D), thus ruling out the possibility of a prevalent mixed identity. Instead, the contour plot and cell proportion by cluster (Figure 5D-E), pointed to a reduction in the intermediate populations in CHIR treated organoids at day 50 and day 100, particularly affecting the cluster 4 at day 100 that is abundant in oRG. This observation was supported by the decreased number of HOPX expressing cells in cluster 4 (Figure 5C), suggesting that in the absence of GSK3 activity, oRG-independent neurogenic trajectories are favored. In order to probe the mechanisms underlying these differences, we harnessed previously defined gene signatures characteristic of cell cycle phases (S, G2M and G1-G0 by subtraction) from single cell datasets (Tirosh et al., 2016). Under the assumption of the cell cycle length being equal in homogeneous populations, we used the proportion of cells expressing these signatures as an estimation of the duration of each cycle phase. By this approach, we found that the cells in intermediate progenitors and early neural progenitors from CHIR-treated organoids have a 15-20% increase G1 phase length at day 100 (Figure. 5F), in agreement with a well-described relationship between G1 lengthening and neurogenesis promotion (Calegari, 2005) and consistent with the higher proportion of mature cells found at the same developmental stage (Figure 5D). Next, to further investigate the relations between populations, we used partition-based graph abstraction (PAGA), as a way to estimate the strength of connectivity across the cells belonging to each cluster, thus inferring transitional links among subpopulations. This approach confirmed a decreased of connections between intermediate progenitor clusters and oRG clusters (yellow) upon GSK3 inhibition, particularly evident at day100 (Figure 5G) along with a global decrease in direct links between the areas corresponding to these subpopulations, corroborating an overall reduction of the intermediate-maturity transition.

Finally, by applying pseudotime independently to all conditions and stages, we identified a second developmental trajectory that becomes apparent by day 100 in controls and corresponds to an oRG identity (Figure 5H). Strikingly, GSK3 inhibition resulted in a trajectory preference towards direct neurogenesis (Figure 5H) which could be reproduced bidirectionally by decomposing trajectories into first and second components (Figure S5B). In agreement, HOPX distribution of expression peaked at day 100 in control organoids at late pseudotime, while remaining stalled in day 100 CHIR-treated organoids (Figure 5H). Together, these results indicate that GSK3 inhibition results in a severe reduction of HOPX expressing cells and potentially oRG-dependent lineages.

## Discussion

This work provides the first dissection of the role of GSK3 activity throughout early to mid-fetal corticogenesis in a human background. Specifically, our integrated analysis of tissue architecture and transcriptional regulation at single cell resolution uncovers a key role of GSK3 activity in cortical morphogenesis, with a surprisingly selective impact on neuronal output, as evidenced by the cell cycle-dependent acceleration of an otherwise uncompromised early neurogenesis vis à vis the strong decrease of oRG production.

An unexpected finding was the mildness of the defect in early neurogenesis in the face of the increased progenitor proliferation and massive morphologic disarray of CHIR-treated organoids. This represents a conspicuous example of the relative disconnect between tissue morphogenesis and transcriptional identity, since in organoids the latter matches frequently with high fidelity the transcriptional milestones of human fetal development even across significant morphological heterogeneity (Camp et al., 2015; Luo et al., 2016; Mariani et al., 2015; Quadrato et al., 2017). This emphasizes the resilience and degree of cell-autonomy of the transcriptional programs involved in neuronal identity. Indeed, the expression levels of stage-wise neuronal maturity determinants as well as the abundancy of TBR1+ cells, showed only marginal differences in either direction at day 50 and basically no differences at day 100. Dissection of single cell populations at day 100 allowed to pinpoint selective differences in specific subsets of mature populations, differentially affecting the expression of gene sets including several neuronal fate and function modulators, highlighting a surprising specificity for GSK3 impact on fate specification.

Depletion of GSK3 activity using pharmacological inhibition or knockdown causes drastic changes in radial glial organization (Yokota et al., 2010). Polarization of radial glia is prerequisite for cortical scaffolding, proper neuronal migration and layering (Shah et al., 2017) and thus alterations of this process results in impaired cortical plate formation (Beattie et al., 2017; Shah et al., 2017). The effects of GSK3 in radial glia organization constitute fast cellular responses triggered by phosphorylation of targets such as CRMP2, MAP1B and CLASP, which in turn effects modifications and structural changes in microtubule cytoskeleton (Hur and Zhou, 2010; Yokota et al., 2010). We did not observe any differences in expression of canonical polarity markers or determinants of radial glia organization (data not shown), thus in the human setting, GSK3 activity has no downstream impact on the regulation of expression of polarity determinants. Rather, the most salient transcriptional global effect was the upregulation of modulators of cell replication and, in particular, the G2M transition emerged as a recurrent target at all stages, indicating that the modulation of the G2M is a persistent feature of GSK3 activity throughout human corticogenesis.

During human corticogenesis, most neurons are not generated directly from apical radial glia, but rather indirectly via IPCs that originate from aRG and are located in the subventricular zone (Bystron et al., 2008; Florio and Huttner, 2014). Depending of the context, depletion of GSK3 may result in either reduced or increased neurogenesis in mice (Kim et al., 2009), indicating that spatiotemporal regulation of GSK3 activity is required for an appropriate transition from the proliferative to the neurogenic phase occurring during brain development. In our chronic setting, starting from a pluripotent state (day 0), GSK3 inhibition caused increased proliferation and polarity defects in NSCs, reflected also in larger organoid size and increased number of neurons. The dramatic increase in transcriptional impact at day 50 compared to day 18, suggests that the early defects in NSCs are amplified in intermediate and committed progenitors, indicating a higher reliance on GSK3 activity at later stages and particularly in IPCs. In agreement, the population breakdown by single cell analysis revealed a decrease of IPCs and in particular oRGs as well as an increase of the duration of IPC G1 phase, which is well documented to promote neurogenesis by accelerating the entrance in a postmitotic state (Calegari, 2005).

Pseudotime trajectories showed that cortical organoids recapitulate the co-existence of an IPC-mediated neurogenic trajectory, visible by day 50, that is juxtaposed to an indirect trajectory reaching an oRG identity by day100, in agreement with a protracted wave of neurogenesis dependent on oRG production, known to be a salient feature of primate corticogenesis and responsible of human cortex lateral expansion (Bershteyn et al., 2017; Florio and Huttner, 2014; Nowakowski et al., 2016). CHIR-treated organoids displayed a strong downregulation of the early onset neural transcripts NEUROD1/2, without affecting NEUROD6 or NEUROG2, underscoring a rather selective impact of GSK3 in neurogenic fates. Likewise, outer radial glia numbers were significantly depleted in CHIR-treated organoids accompanied by a depletion of the early astrocytic markers SPARCL1 (Sloan et al., 2017; Zhang et al., 2016), suggesting a potential impact in oRG-mediated astrogenesis. The disappearance of the indirect neurogenic trajectory at day100 upon CHIR treatment points to a key role of GSK3 in the establishment of IPC/oRG populations. Our results indicate that in humans, GSK3 activity is not essential for early RG-dependent neurogenesis, with only selective impact on specific fates, is pivotal for later oRG generation and consequently oRG-dependent lineages.

## Author contributions

Organoid differentiation and sample collection for the different experiments ALT, ST, NC, ML and BMC. CEV and CC bioinformatic analysis. ALT, ST and NC single cell experimental procedures. ALT and ML organoid stainings and growth curve, ALT and PLR quantifications and statistical analysis. PC, RC and EC neural induction experiments in 2D. MTR performed proliferation assay in 2D. MDS, ILN, GM, SP and MP contributed with funding and expertise. ALT wrote and the manuscript with contributions from all the authors. GT conceived, designed and supervised the study.

## Acknowledgements

This work was supported by the Associazione Italiana per la Ricerca sul Cancro (AIRC) (IG 2014-2018 to G.T.); EPIGEN Flagship Project of the Italian National Research Council (CNR) (to G.T., S.P. and M.P.); the European Research Council (DISEASEAVATARS to G.T.); Fondazione Cariplo (grant 2017-0886 to A.L.T.); Fondazione Italiana per la Ricerca sul Cancro (FIRC) (to P.L.R.); the AIRC grant n° IG2016-ID18575 (to M.P.) and the ERC Consolidator Grant n° 617978 (to M.P.) We are grateful to Pierre-Luc Germain for providing scripts that facilitated this work. To Andreas Püschel for critical comments to the introduction and to Federica Pisati from tissue processing facility.

